# The Baldwin effect reloaded: Intermediate levels of phenotypic plasticity favor evolutionary rescue

**DOI:** 10.1101/2025.01.17.633614

**Authors:** Amaury Lambert, Guillaume Achaz, Arnaud Le Rouzic, Laurent Loison

**Affiliations:** Institute of Biology of ENS (IBENS), École Normale Supérieure (ENS-PSL), CNRS UMR 8197, INSERM U1024, PSL University, Paris, France; Center for Interdisciplinary Research in Biology (CIRB), Collège de France, CNRS UMR 7241, INSERM U1050, PSL University, Paris, France; Université Paris-Cité, Paris, France; Laboratoire Évolution, Génomes, Comportement, Écologie (EGCE), CNRS UMR 9191, IRD, Université Paris-Saclay, Gif-sur-Yvette, France; Laboratoire Sciences, Philosophie, Histoire (SPHERE), CNRS UMR 7219, Université Paris-Cité, Paris, France

**Keywords:** population genetics, demography, stochastic model, adaptation, extinction, environmental variability, conservation, mathematical evolution

## Abstract

Since the late 1890s and until today, how phenotypic plasticity interacts with genetic adaptation is a debated issue. Proponents of a positive causal role of phenotypic plasticity –James M. Bald-win in the first place– supported the view that, in altered environmental conditions, phenotypic plasticity is a key factor allowing a population to avoid extinction and then genetic evolution to catch up (“Original Baldwin Effect”, thereafter OBE). Opponents, like for instance Ernst Mayr, regularly pointed out that phenotypic plasticity, by masking genetic variation, slows gene-level evolution (“Mayr Effect”, thereafter ME). For decades this opposition remained only verbal and qualitative. To resolve it, we propose here a stochastic model that, following Baldwin’s intuitive take, combines the minimal number of ingredients to account for extinction, selection, mutation and plasticity. We study evolutionary rescue of the population (arrival and invasion of an adaptive genetic mutant) in the altered environment for different values of phenotypic plasticity, here quantified as the probability *p* that the maladapted genotype develops into the adapted phenotype. Our claim is that OBE can be a genuine evolutionary mechanism, depending on the level of phenotypic plasticity with respect to a threshold value *p*^*⋆*^: when *p < p*^*⋆*^, increasing *p* promotes evolutionary rescue by delaying extinction (“Strong” OBE); when *p > p*^*⋆*^, plasticity sustains population survival and increasing *p* has two antagonistic effects: to accelerate adaptation by increasing the supply of adaptive mutants (“Weak” OBE, intermediate values of *p*), and to slow down adaptation by decreasing their fitness advantage (ME, high values of *p*).

## Introduction

Since the initial publication of James M. Baldwin in the *American Naturalist* (1896), the way plasticity interacts with adaptive evolution has no less than haunted the scientific literature for more than a century. Terms like “organic selection”, “coincident selection”, “post-adaptation”, “Baldwin effect”, “stabilizing selection”, “genetic assimilation” evidently show that a clear consensus has never been reached. The related issues of the definition, the causal mechanism at stake, and the empirical basis of these possible evolutionary processes have remained open questions ever since.

This work does not aim to explore all these evolutionary models, which are sometimes very different from one another (Loison 2025). In this article, we focus exclusively on the ecological consequences of phenotypic plasticity, as a property of living systems that enables certain populations to survive drastic variation in their environment. In doing so, this work is in line with Baldwin’s own original intuitions, while the possible causal role of plasticity for evolution has been positively considered again over the last two decades or so (West-Eberhard 2003; Weber and Depew 2003; Pigliucci 2010). Our work takes place within this debate and intends to contribute to its renewal.

The numerous ambiguities attached to the evolutionary scope of phenotypic plasticity require at least a double effort. First, we need to understand how this class of concepts (like “organic selection” and then the “Baldwin effect”) were understood throughout the history of science, in order to grasp their more central aspects. The first section of this article is therefore a historical investigation, in which we will see that at least two different “effects” of plasticity on genetic evolution were conceptualized. The first, termed here “Original Baldwin Effect” (OBE) assumes that plasticity is necessary to avoid extinction, which will then enable genetic evolution to catch up. The second, “Mayr Effect” (ME), partly opposes this hypothesis on the ground that plasticity masks fitness differences and thus is an obstacle to gene-level evolution.

We then propose to resolve this debate by quantifying in the same framework the effects of selection, mutation, plasticity and demography, in order to understand to what extent OBE and/or ME can play an actual role (positive or negative) in adaptive evolution. We introduce a model designed to be as general as possible and include all required ingredients to account for genetic drift, variable population size, variable environment, competition, mutation and plasticity, but remains simple. In this model, a crucial role is played by the value *p* of phenotypic plasticity, defined here as the probability that a carrier of the wild genotype, assumed to be maladaptive in the altered environment, develops into an adaptive phenotype.

We study the likelihood of arrival and invasion of a genetically adapted mutant, a phenomenon nowadays termed “evolutionary rescue” and the waiting time before evolutionary rescue as a function of *p*. Our main result is that this time is minimum at an intermediate value of *p* – thereby reconciling OBE and ME: at low values of *p*, extinction occurs before the first successful mutant can arise, preventing genetic adaptation (i.e. OBE is inefficient); whereas at high values of *p*, the selective advantage of the adaptive genotype vanishes, reducing the probability of invasion of its carriers (i.e. ME progressively takes the lead).

We also distinguish between a strong version of OBE, where the population benefits from a non-trivial join effect of mutation and plasticity and a weak version of OBE, where plasticity alone is powerful enough to save the population and thus allows for subsequent genetic adaptation with certainty (provided adaptation mutation rate is nonzero). We use a mathematical framework of large equilibrium population size and small mutation rate enabling explicit calculations, in particular of the probability and distribution of time rescue, as well as rescue from standing variation. These mathematical results are supported by extensive simulations.

### Which Baldwin effect? A tale of two (opposite) effects

Scientific concepts are fluid entities that evolve through time. Their definitions, mathematical formalism or experimental basis are subjected to change. It so happens that some conceptual content can be lost in the course of history, as is the case, we believe, for the Baldwin effect and related concepts. This history is mainly divided into two quite distinct parts. During the period 1896-1953, several ethologists, zoologists and geneticists questioned the possible positive role of phenotypic plasticity in adaptive evolution. These biologists all rejected the possibility of a *direct* role, through a physiological mechanism of inheritance of acquired characters (Lamarckism). They believed that phenotypic plasticity is an evolutionary causal factor (and not only an evolutionary product) because, in altered environments, it can allow the transient survival of the population and, later on, the fixation of adaptive variants. Such an idea is well captured by the recent expression of a “breathing space” (Godfrey-Smith 2003). This central aspect can be found both around 1900, when the speculative hypothesis termed “organic selection” was formed for the first time, and forty years later, when this evolutionary mechanism attracted a renewed (yet limited) interest at the time of the rise of the Modern Synthesis.

From 1896 onwards, James M. Baldwin himself always highlighted that the evolutionary consequence of plasticity was to “keep single organisms alive” (Baldwin 1896) and that this will “give the species time to acquire the congenital mechanism [genetic mutations] for the same functions [adaptive phenotype]” (Baldwin 1902). This aspect was also central for the other co-founders of this “new factor in evolution” (the title of Baldwin’s seminal paper). Ethologist Conway Lloyd Morgan spoke of “escaping elimination in the life struggle”, so that “the organism waits, so to speak, for a further congenital [hereditary] variation” (Morgan 1896). For paleontologist Henry F. Osborn, plasticity “enables animals and plants to survive very critical changes in their environments” (Osborn 1897).

After a 30-year eclipse (roughly 1905-1935), “organic selection” started to be reworked in various theoretical contexts under different names (“coincident selection”, “post-adaptation”, etc.). Here again, the idea that plasticity provides a “breathing space” in altered environments was commonly shared and put at the forefront of the mechanism (Hovasse 1943). The few attempts to include this still unlabeled mechanism within the nascent Modern Synthesis framework made at first no exception. Already in 1942, in his seminal book *Evolution, the Modern Synthesis*, ethologist and zoologist Julian Huxley emphasized the fact that phenotypic variations “may serve as the first step in evolutionary change, not by becoming impressed upon the germ-plasm [Lamarckism], but by *holding the strain* in an environment where mutations tending in the same direction will be selected and incorporated into the constitution” (Huxley 1942, p. 304, our emphasis).

During the first half of this history, the effect of plasticity on adaptive evolution was thus conceived as positive by the supporters of organic selection and the like: plasticity enabled the transient survival of a population facing a new environmental challenge, allowing the standard mutation/selection process to take over. We term this positive effect of plasticity, expressed verbally during the period 1896-1953 (including by Baldwin himself), the “Original Baldwin Effect” (OBE).

The turning point was George G. Simpson’s 1953 paper where he proposed to call this set of mechanisms based on phenotypic plasticity the “Baldwin effect”. The term “Baldwin principle” had already been proposed a few years earlier (Hovasse 1950), but Simpson preferred “effect” to “principle”, as the evolutionary result seemed to him, quite rightly, to be the conjunction of a series of phenomena that could add up their effects. As a result, he tried to summarize this causal series in the form of a 3-step process (Simpson 1953):

1. Individual organisms interact with the environment in such a way as systematically to produce in them behavioral, physiological, or structural modifications that are not hereditary as such but that are advantageous for survival, i.e., are adaptive for the individuals having them
2. There occur in the population genetic factors producing hereditary characteristics similar to the individual modification referred to in (1), or having the same sorts of adaptive advantages.
3. The genetic factors of (2) are favored by natural selection and tend to spread in the population over the course of generations. The net result is that adaptation originally individual and non-hereditary becomes hereditary.

Remarkably, the issue of extinction is absent from Simpson’s verbal characterization. Moreover, this looks more like a description than a causal explanation and, in Simpson’s take, it is difficult to understand why such a process should take place at all. This conceptual impoverishment was to be found in many later works, notably those of his colleague Ernst Mayr. On this basis, Mayr strongly doubted the very possibility of plasticity-first evolution on the ground that “if the phenotype is highly plastic, the selection pressure may actually be reduced because there is no selective advantage in changing the genotype when an individual can adjust itself phenotypically to a current condition” (Mayr 1970). In order to avoid confusion, we term this negative effect of plasticity on genetic evolution the “Mayr Effect” (ME).

From this time on, within Simpson’s framework, most of the debates were only about whether adaptive plasticity – “learning” in most cases – accelerates (by channeling the exploration of the phenotypic space) or decelerates (by masking genetic variation) the evolutionary process (Mayley et al. 1997; Sznajder et al. 2012). Hinton and Nowlan’s 1987 model was especially influential in this respect and was discussed during years, assuming that with no phenotypic plasticity, genetic adaptation would eventually occur, yet more slowly (Maynard Smith 1987; Belew 1989; Gruau and Whitley 1993; Mayley et al. 1997; Weber and Depew 2003; Le 2019).

In the past ten years or so, OBE, i.e. the idea that “plasticity is buying time” (Pennisi 2018; Diamond and Martin 2021) has resurfaced. It is again argued that without plasticity, population will go extinct before an adaptive variant can be selected. Based on what history tells us, we believe that the core theoretical issue is thus not the speed of adaptation, slow versus fast (Paenke et al. 2007), but, more fundamentally, to decide between the survival effect of plasticity (OBE) and the masking effect of plasticity (ME): phenotypic plasticity can potentiate population survival (OBE) but also prevent genetic evolution (ME, see Price et al. 2003).

Which effect will take the lead? This question cannot be solved on a conceptual ground only, and needs a quantitative model that addresses both aspects at the same time (Ashander et al. 2016). In such a model, a key ingredient has to be the population size, an aspect often neglected so far. This verbal characterization grounds our theoretical model in order to quantitatively compare the efficiency of two different and opposite effects of plasticity on genetic evolution, namely OBE and ME.

## Model

### Plasticity in population genetic models

Whether or not phenotypic plasticity and its evolutionary consequences are fully integrated into the contemporary version of the Modern Synthesis remains debated (Pigliucci and Müller 2010; Laland et al. 2014; Futuyma 2021). All evolutionary biologists, including Darwin himself, have acknowledged the existence of plasticity, but few have explicitly merged it into the mainstream evolutionary thinking. Unlike most genetic mechanisms that have been progressively added into the formal population genetics corpus during the 20th century (including dominance, epistasis, linkage disequilibrium, mutations, selfing, assortative mating, pleiotropy, maternal effects, genetic drift, or population structure), formal models of phenotypic plasticity are rather recent, and are characterized by a wide diversity of theoretical frameworks.

Historically, the first models capturing phenotypic plasticity in quantitative genetics were justified by the need to account for different performances of plant or animal strains across different environments (reviewed in Lynch and Walsh 1998). Based on the traditional decomposition of variances (Cockerham 1963), it is possible to estimate an “environmental” effect on the phenotype (plasticity), and a genetic × environment interaction reflecting the genetic variation in plasticity. However, if such a statistical description of plasticity can be used to predict the effect of plasticity on phenotypic evolution, it remains unable to predict how plasticity itself would evolve as a consequence of selection.

To do so, theoretical evolutionary genetic models focus on a heritable character that can either be an environment-dependent phenotype (Via and Lande 1985), the norm of reaction (e.g., slope and elevation for linear reaction norms) (Gavrilets and Scheiner 1993) or the (environment-dependent) fitness (Orzack 1985; Moran 1992; Wu et al. 2014; Kang and Park 2018), as we do here. More complex genetic architectures, featuring pleiotropy and epistasis, can be addressed by numerical models of arbitrary genotype-phenotype complexity (Bull 1999; Suzuki and Arita 2007), or by gene regulatory networks (Masel 2004; Kaneko 2012; Brun-Usan et al. 2021; Burban et al. 2022). Population genetic models, based on a small number of genetic factors, have also handled plasticity in many different ways: plasticity can be represented as a set of environments in which a genotype displays maximal fitness (Ancel 1999, 2000), as a capacity to generate random phenotypic variation (Majic et al. 2021), or as a set of behavioral decision rules in pairwise interactions among individuals (Suzuki and Arita 2004).

In the theoretical literature, references to the Baldwin effect are neither consistent nor systematic. Depending on the context, the Baldwin effect may refer to the evolution of learning (Suzuki and Arita 2004), to the accelerating effect of plasticity on evolution (Hinton and Nowlan 1987; Ancel 2000), or as a general synonym for genetic assimilation (Waddington 1953*b*).

For example, in computer science, *plasticity* and *learning* are often intimately intermingled. In the landmark article by Hinton and Nowlan (1987), a fixed size population of genotypes is evolved. Genomes have 20 loci with three possible alleles: ‘0’, ‘1’ or ‘?’. The fitness landscape is entirely flat, except for a unique peak at all ‘1’s (“111…111”). Each initial genotype has ten loci randomly set to ‘1’ or ‘0’ (genetically determined) and ten loci set to ‘?’ (plastic loci). During their lifetime, genomes search the peak by setting the ‘?’ loci to 0 or 1. The fitness of a genotype is evaluated by measuring how long it takes to find the peak. As an effect of selection and recombination (and mutations in subsequent studies), surviving genotypes only carry ‘1’ and ‘?’ alleles, and the plastic ‘?’ alleles are progressively replaced by fitter, genetically determined ‘1’ alleles. Following, a number of studies have explored the interaction between learning and evolution (Turney et al. 1997). Among them, a study where the landscape includes a second unique lower peak with fifteen ‘1’s (Mills and Watson 2006). The Baldwin effect is then discussed in the context of finding and then escaping this local peak and more generally how learning can facilitate crossing valleys of fitness. In these algorithms, population size is held constant so that the risk of extinction is zero.

Most of the theory about evolutionary rescue focuses on non-plastic traits (Gomulkiewicz and Holt 1995; Bürger and Lynch 1995; Chevin et al. 2017; Orr and Unckless 2014; Björklund et al. 2009), and most of the theory about the evolution of plastic traits consider constant population sizes (Lande 2009; Kang and Park 2018). The effect of plasticity on the population size has been explored recently in a quantitative genetics framework; plasticity can be proven to decrease the drop in population size after an environmental change in a deterministic setting (Chevin and Lande 2009; Greenspoon and Spencer 2021), and to decrease the probability of extinction in a stochastic setting (Ashander et al. 2016). These models confirmed that theory predicts the evolution of plasticity during evolutionary rescue, associated with an increase in the genetic variance of the trait. Plasticity is also expected to be lost over long evolutionary time when sub-optimal (for instance, when environmental cues are imperfect) (Bell 2017). The potential for phenotypic plasticity to buy time and allow for evolutionary rescue has sometimes been noticed in simulations (Reed et al. 2010; Carja and Plotkin 2019), but these observations were not connected with historical concepts, and whether this should be interpreted as a theoretical support for the Baldwin effect remains unclear. Here, we propose to focus explicitly on the conditions in which plasticity and genetic adaptation could realistically interact and affect the evolution of populations.

### Our model

The model here proposed is a stochastic model of population genetics featuring the minimal number of ingredients to model genetic drift, variable population size, competition, mutation and phenotypic plasticity. This setting is similar to traditional evolutionary rescue models (Orr and Unckless 2014) with two major differences: 1) here, population size is mediated by ecological interactions rather than fixed a priori, and most notably, is allowed to go to 0 (extinction), and 2) our model includes varying degrees of phenotypic plasticity. We use the same kind of modeling assumptions as Champagnat (2006), parameterized by a scaling parameter *K* thought to be large, where mutations are rare (mutation probability of the order of 1/*K*) and resident population is large (population size of the order of *K*), as an effect of weak competition (pairwise competition death rate of the order of 1/*K*).

We consider a population of asexual, haploid individuals living in an environment with two possible states, named *E*_0_ and *E*_1_.

### Genotypes and phenotypes

Each individual is endowed with a genotype with two alleles *A* and *B*, and with a phenotype that can take two values, *α* and *β*.

We think of phenotype *α* (resp. phenotype *β*) as the natural phenotype of carriers of allele *A* (resp. allele *B*). We also think of *α*-individuals as well adapted to *E*_0_ and maladapted to *E*_1_ and conversely of *β*-individuals as maladapted to *E*_0_, but well adapted to *E*_1_.

### Phenotypic plasticity

We assume that in *E*_1_, each individual carrying allele *A* develops into phenotype *α* with probability 1 − *p* and into phenotype *β* with probability *p*, independently of everything else. The parameter *p* quantifies the ability of *A*-individuals to adapt to *E*_1_ despite carrying the maladapted allele, and is thus the natural measure of phenotypic plasticity in our setting.

We could easily extend this assumption of phenotypic plasticity to other environments and/or genotypes, without qualitatively changing our results, but we refrain from doing so, for the sake of simplicity. Specifically, all *A*-individuals develop into phenotype *α* in *E*_0_, and in both environments, *B*-individuals always develop into phenotype *β*.

### Genetic mutations

We assume that genotypes can mutate and we denote by *µ*_*i*_ the probability of mutation at birth from a mother carrying allele *i*. For example, each time an *A*-individual gives birth, the newborn carries allele *A* with probability 1 − *µ*_*A*_ and allele *B* with probability *µ*_*A*_. As is standard in population genetics, we set *θ*_*A*_ := *Kµ*_*A*_ and *θ*_*B*_ := *Kµ*_*B*_ the effective mutation rates, where *K* is a parameter scaling similarly as population size, intended to be large.

### Population dynamics

We use a classic Markov model of population dynamics featuring density-dependent competition. We assume that in *E*_*k*_, *k* = 0, 1, when the current population size is *n*, each individual of phenotype *j*, regardless of its genotype, gives birth at rate *b*_*j*|*k*_ and dies at rate *d*_*j*|*k*_ + *c*_*j*|*k*_(*n* − 1)/*K*. We write *r*_*j*|*k*_ := *b*_*j*|*k*_ − *d*_*j*|*k*_ for their natural growth rate when *n* « *K*.

### Assumptions

To implement the idea that *α*-individuals are better adapted than *β*-individuals to *E*_0_, we assume that

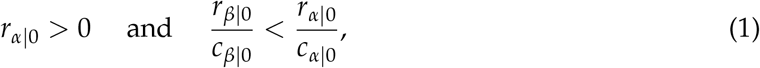

which holds for example if *r*_*β*|0_ *<* 0.

To implement the idea that in *E*_1_, *β*-individuals are adaptive but *α*-individuals are maladaptive, we assume that

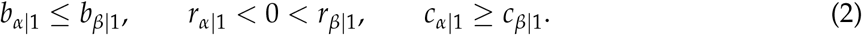

## Results

### Large population, rare mutations heuristic

Here we give a heuristic description of the behavior of the model when *K* is large. Recall that by assumption, the *per capita* death rates due to competition scale like *n*/*K* (where *n* denotes the current population size) and the *per capita* mutation rate scales like 1/*K*. As a consequence, 1) it is only when the population size is of order *K* that individuals feel the competition, 2) the total population size is of order *K* at equilibrium, and 3) it is only when the total population is of order *K* that the first mutations begin to arise.

Let 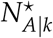 be the equilibrium size of the *A*-population in *E*_*k*_, *k* = 0, 1, in the absence of *B*-carriers. In *E*_0_, at equilibrium, births and deaths of *A*-carriers are balanced so that 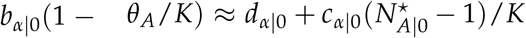. This yields 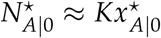, where (see Appendix for a rigorous derivation)

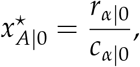

which is positive because *r*_*α*|0_ *>* 0 by Assumption (1). In *E*_1_, we define

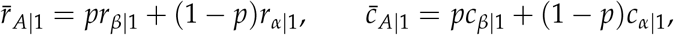

which are respectively the average growth rate and the average competition rate of *A*-carriers in *E*_1_. By Assumption (2), *r*_*α*|1_ *<* 0 but *r*_*β*|1_ *>* 0, so that 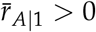 whenever *p > p*^*⋆*^, where

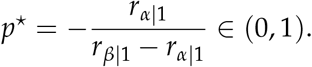

Then by the same reasoning as previously (again see Appendix), 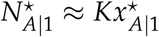, where

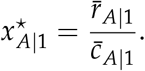

When the *A*-population sits at its equilibrium size 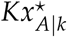, it fuels the *B*-population with mutations occurring at rate 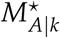 equal to the product of 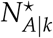 with the *per capita* rate of births-with-mutation. In *E*_0_,

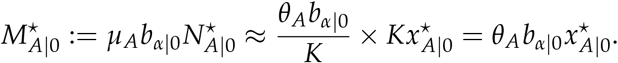

Similarly, defining 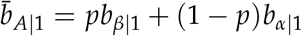,

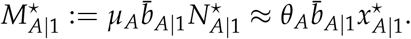

As a conclusion,

- In *E*_0_, the *A*-population outcompetes the *B*-population; the former sits at large equilibrium Size 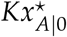 and the latter fluctuates at low numbers under mutation-selection equilibrium;
- In *E*_1_, the *B*-population outcompetes the *A*-population but the *B*’s are assumed to be initially absent from the population. We are thus interested in the time it takes for a *B*-mutant to arise and take over (evolutionary rescue).

From now on, we denote by *N*_*i*_(*t*) the number of individuals carrying allele *i* at time *t*.

### First phase – equilibrium in initial environment

In *E*_0_, the *A*-population outcompetes the *B*-population by Assumption (1), so regardless of initial conditions, *N*_*A*_(*t*) converges to the stable state 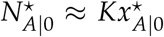, see Figure 1 (and Appendix for a rigorous analysis of this transient phase).

**Figure 1:**
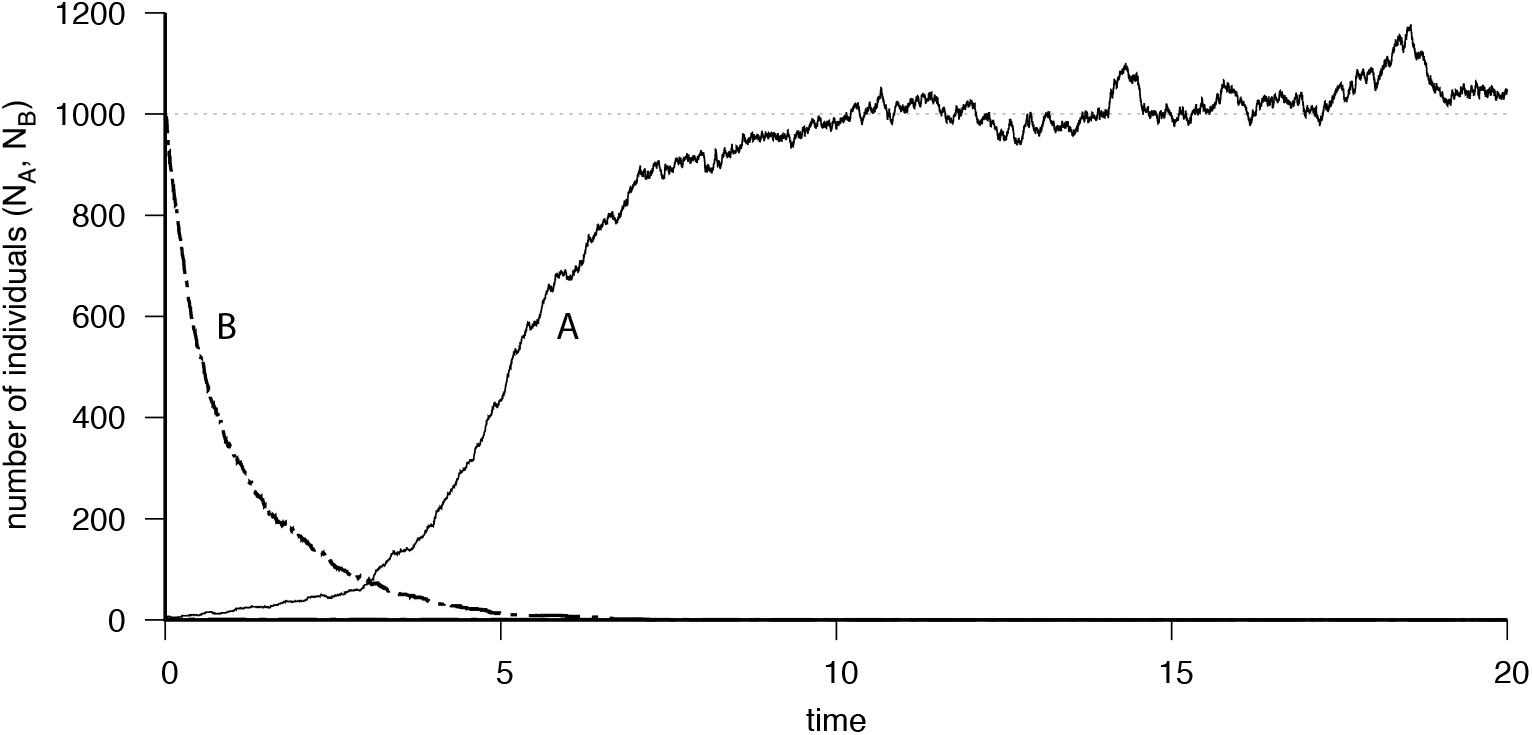
In *E*_0_, genotype *A* outcompetes genotype. *B*. Parameter values: *N*_*A*_(0) = 5, *N*_*B*_(0) = 1000, *b*_*α*|0_ = 2, *d*_*α*|0_ = 1 (*α* is supercritical), *b*_*β*|0_ = 0.5, *d*_*β*|0_ = 1 (*β* is subcritical), *c*_*α*|0_ = *c*_*β*|0_ = 1, *K* = 1000, so that 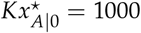 (equilibrium number of genotypes *A*).

In the presence of mutations, this equilibrium is only slightly perturbed by the sporadic presence of *B*’s segregating at low numbers. Indeed, the *per capita* birth rate of *B*’s is *b*_*β*|0_(1 − *θ*_*B*_/*K*) ≈ *b*_*β*|0_ and their total *per capita* death rate is

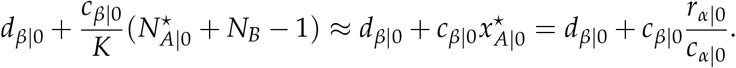

This *per capita* death rate is roughly constant and smaller than *b*_*β*|0_ because *r*_*β*|0_ *< r*_*α*|0_*c*_*β*|0_/*c*_*α*|0_ by Assumption (1). Therefore, *de novo B*-mutants are introduced at rate approximately equal to 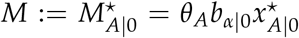 and each of them starts a subcritical branching process with birth rate approximately equal to *B* := *b*_*β*|0_ and death rate approximately equal to 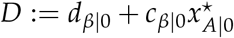.

This can be seen on simulations in Figure 2a, where the *A*-population oscillates around 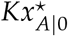 while the *B*-population experiences repeated resurgences, each initiated by the introduction of a mutant and followed by a cycle of births and deaths quickly fading away.

**Figure 2:**
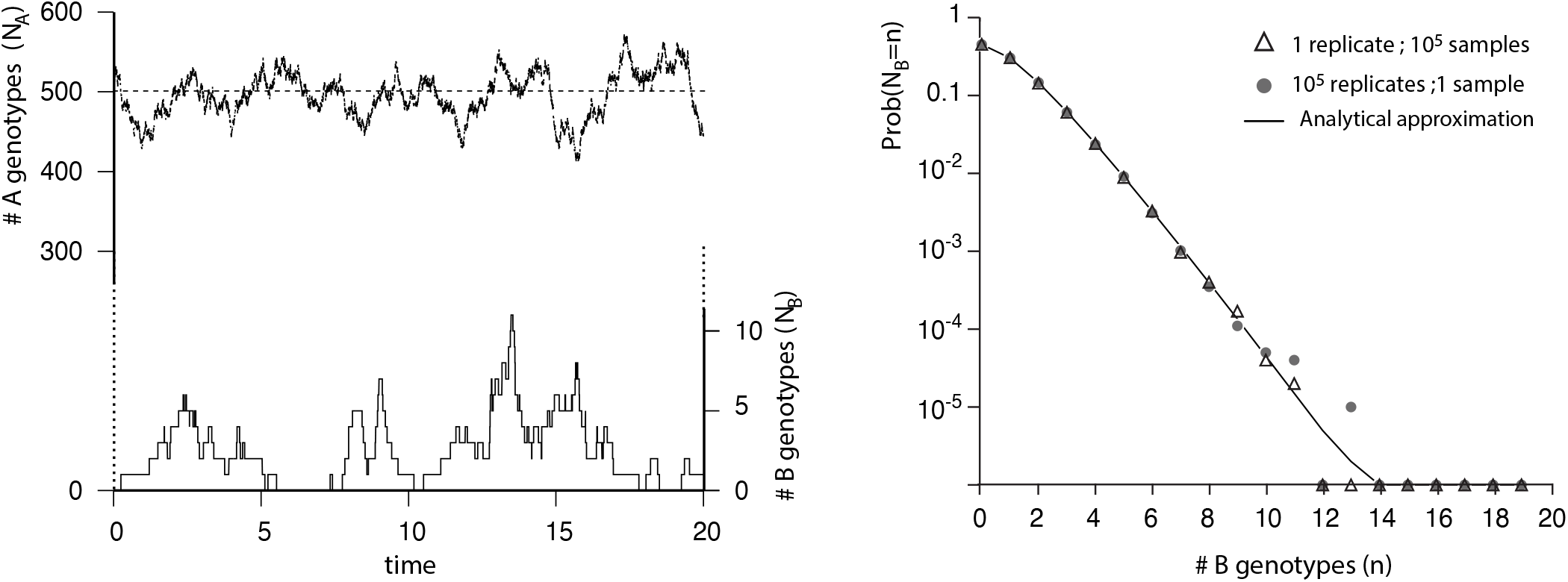
In *E*_0_, genotype *B* fluctuates at low frequency. Left panel: Simulation showing the coexistence through time between the *A*-population, whose size fluctuates around 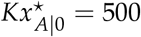, and the *B*-population which has small, random equilibrium size 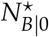 (mind the different scales); right panel: empirical distribution of 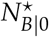 and its analytical approximation. Triangles are 10^5^ samples of the same process, whereas disks corresponds to a single sample of 10^5^ independent replicates. Parameter values: *N*_*A*_(0) = 500, *N*_*B*_(0) = 0, *b*_*α*|0_ = 2, *d*_*α*|0_ = 1 (*α* is supercritical), *b*_*β*|0_ = 0.45, *d*_*β*|0_ = 0.5 (*β* is mildly subcritical), *θ*_*A*_ = 0.5, *c*_*α*|0_ = *c*_*β*|0_ = 1, *K* = 500.

We will write 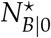 for the (random) size of the *B*-population at this mutation-selection equilibrium. Figure 2b shows the empirical distribution of 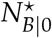, compared to its analytical approximation (see Appendix)

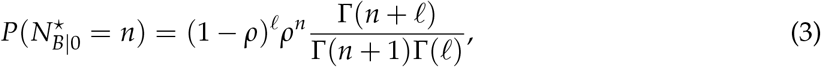

where *ρ* := *B*/*D, 𝓁* := *M*/*B* and Γ is the standard Gamma function.

### Second phase – change of environment

We now assume that at time 0, the population sits at its constant *E*_0_ equilibrium namely 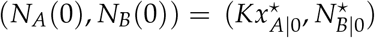, and that the environment suddenly changes from *E*_0_ to *E*_1_.

We are interested in the first time *T* when a *successful B*-mutant arises, where a successful mutant is a mutant whose descendance escapes stochastic extinction and takes over, i.e., grows to its large size equilibrium (order of *K*). We call *T* the *rescue time*.

Note that there is a positive probability that *T* = 0, namely that one of the few *B*’s segregating at mutation-selection *E*_0_ equilibrium, will be successful (adaptation from standing variation, see Appendix). From now on we discard this possibility and study the time *T >* 0 of arrival of the first successful *de novo B*-mutant.

Recall that by Assumption (2), *r*_*α*|1_ *<* 0 *< r*_*β*|1_ and *p*^*⋆*^ = (−*r*_*α*|1_)/(*r*_*β*|1_ − *r*_*α*|1_) is the value of phenotypic plasticity *p* for which the average growth rate of *A*-carriers 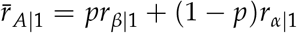 is equal to 0. Since *B*-carriers are assumed to be initially absent,

- For *p* ≤ *p*^*⋆*^, we have 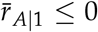 so that *N*_*A*_(*t*) decreases with *t* and eventually goes to 0. The *A*-population becomes extinct, but there is a positive probability that evolutionary rescue occurs before extinction (see Appendix). Increasing *p* delays extinction and thus increases the probability of rescue (see Figure 3, top two rows). We call this effect the Strong OBE;
- For *p > p*^*⋆*^, we have 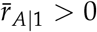 so that *N*_*A*_(*t*) converges to its equilibrium size 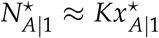 Plasticity is strong enough to sustain the *A*-population at large equilibrium size 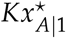 so that rescue sooner or later occurs (*T <* ∞ with probability equal to 1). We call this effect the Weak OBE (see Figure 3, bottom row). However, we will see in the next paragraph that the expectation of *T* has a non-monotonic dependence upon *p*.

**Figure 3:**
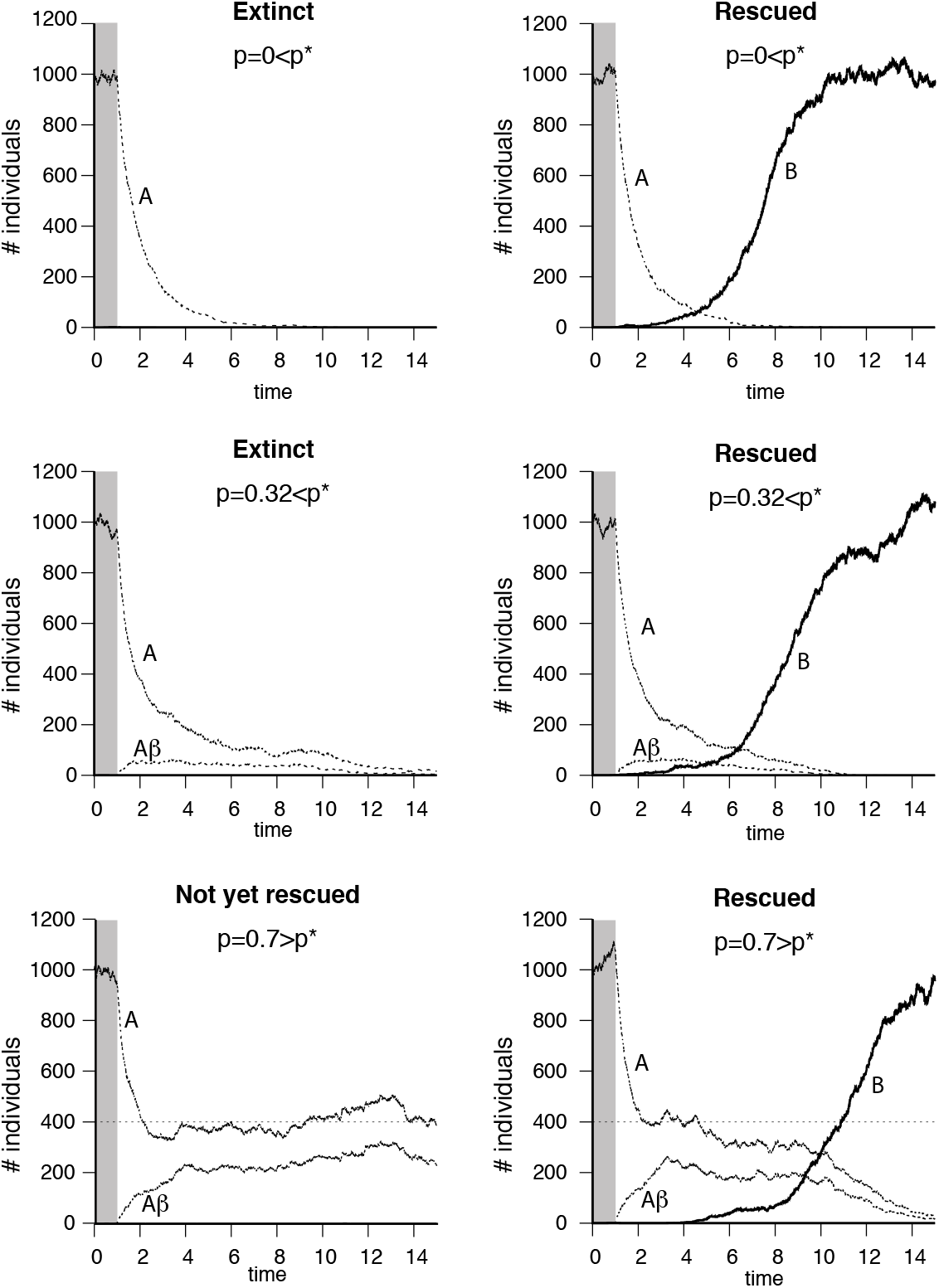
The Original Baldwin Effect (rescue in *E*_1_ by genotype B), depending on plasticity *p*. Gray areas span 1 time unit in *E*_0_ before the transition to *E*_1_. Parameter values: *N*_*A*_(0) = 500, *b*_*α*|0_ = 2, *b*_*α*|1_ = 0.5, *d*_*α*|0_ = *d*_*α*|1_ = 1 (*α* is supercritical in *E*_0_ but subcritical in *E*_1_), *N*_*B*_(0) = 0, *b*_*β*|1_ = 2, *d*_*β*|1_ = 0.5 (*β* is supercritical in *E*_1_), *θ*_*A*_ = 0.5, *c*_*α*|1_ = *c*_*β*|1_ = 1, *K* = 1000. The *A*-population is sustained by plasticity as soon as *p* ≥ *p*^⋆^ = 1/3. Top panel, no plasticity (*p* = 0): the *A*-population goes extinct rapidly (left) but can be rescued by a very lucky *B*-mutant whose descendance shown by bold line takes over (right); central panel, weak plasticity (*p* = 0.32 *< p*^⋆^): extinction is slower thanks to the presence of plastic *Aβ*-individuals whose population size is shown by dashed line (left), which buys time and increases the probability that mutants arise before extinction (Strong Original Baldwin Effect, right); bottom panel (high plasticity, *p* = 0.7 *> p*^⋆^): the population is steadily sustained by plasticity (large subpopulation of plastic *Aβ* individuals, left) so that sooner or later, rescue occurs thanks to the arrival of *B*-mutants who later outcompete the *A*’s (Weak Original Baldwin Effect, right).

### Quantifying the rate of adaptation

Here we assume that *p > p*^*⋆*^. Then the rescue time *T* is always finite and we use the inverse of its expectation as a natural measure of the speed of adaptation/evolutionary rescue. More specifically, we set

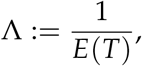

that we call the *rate of adaptation*.

To compute Λ, we make the simplifying assumption (see Appendix for a generic treatment) that we can neglect the transient phase during which *N*_*A*_ goes from its equilibrium 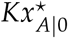 in *E*_0_ at time 0 to its (smaller) equilibrium 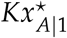 in *E*_1_. Then, while *N*_*B*_ « *K, N*_*A*_ remains approximately constant to 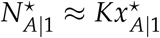 and

- *De novo B*-mutants arise at the times of a Poisson process with intensity 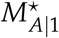;
- The descendance of each *B*-mutant follows a supercritical birth-death process with birth rate *b*_*β*|1_ (1 − *µ*_*B*_) ≈ *b*_*β*|1_ and death rate 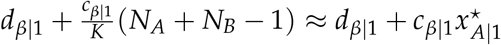;
- The probability for a *B*-mutant to take over is roughly the probability of non-extinction of this supercritical birth-death process starting from one individual, that we denote 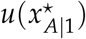, known to be

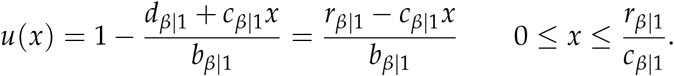

The arrival times of successful *B*-mutants is a subset of the arrival times of *B*-mutants. The former is obtained by keeping each point of the latter independently with probability 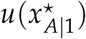. By the thinning property of Poisson point processes, successful *B*-mutants occur according to a Poisson process with intensity 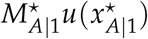. Since *T* is the minimum of this set, it follows the exponential distribution with parameter 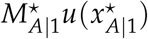. The parameter of an exponential random variable is also the inverse of its expectation, which by definition here is the rate of adaptation:

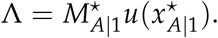

This expression plainly displays the antagonistic effects of *p* on the rate of adaptation. Recall from Assumption (2) that *b*_*α*|1_ ≤ *b*_*β*|1_, *r*_*α*|1_ *< r*_*β*|1_ and *c*_*α*|1_ ≥ *c*_*β*|1_. As a consequence, because increasing *p* increases the probability of displaying phenotype *β* rather than phenotype *α*, increasing *p* increases 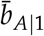, increases 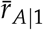, decreases 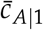, and so increases 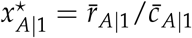. As a consequence (see Figure 4), increasing *p*

**Figure 4:**
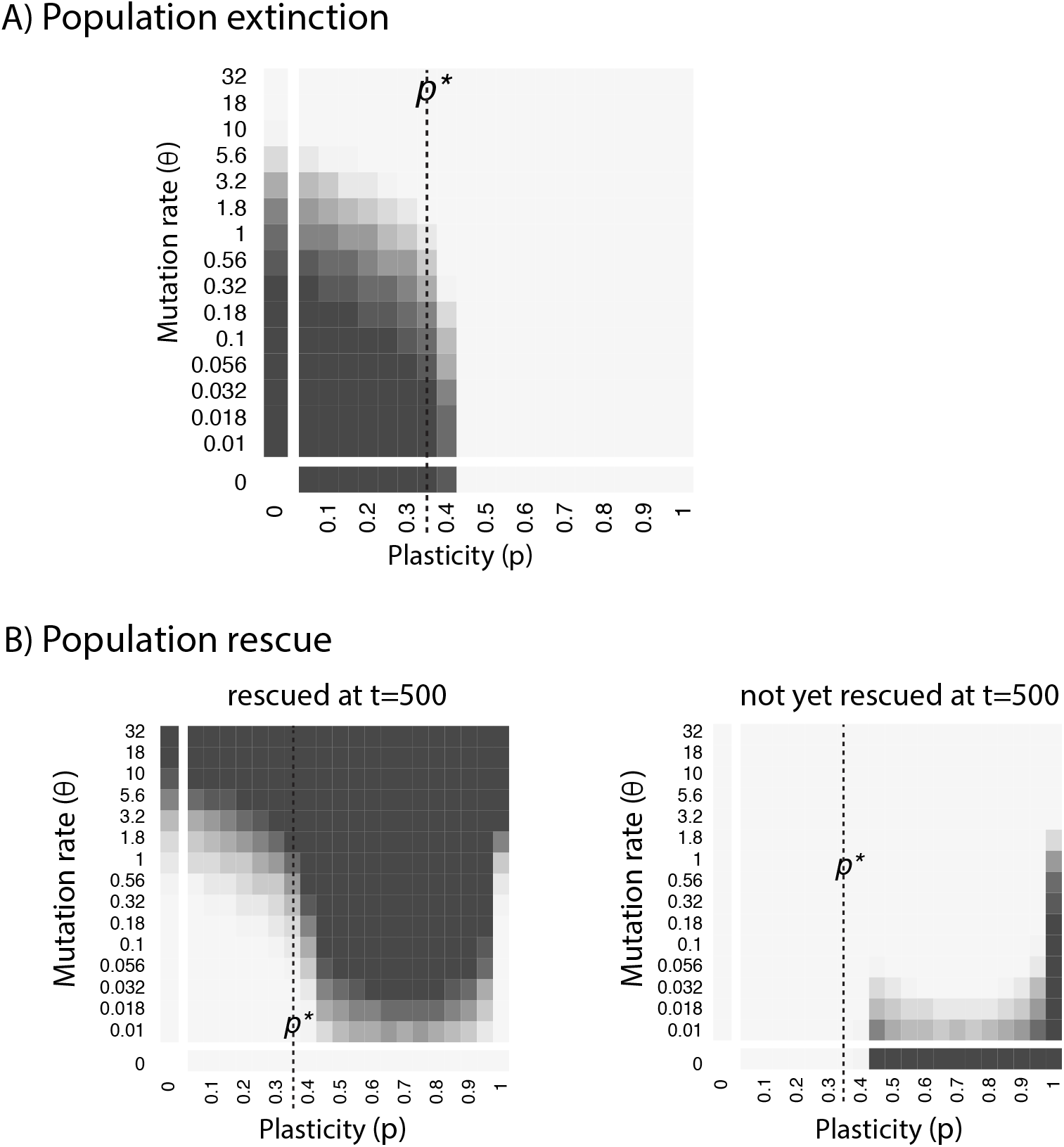
Partitioning the mutation rate *vs* plasticity parameter space. Each panel shows the *θ vs p* plane, where *θ* = *Kµ*_*A*_ is the rescaled rate of mutation from genotype *A* to genotype *B* and *p* is the probability for *A*-carriers to display phenotype *β* in *E*_1_ (phenotypic plasticity). Light gray indicates 0% of the 10^3^ replicates, whereas black indicates 100% of the replicates. Parameter values are identical to Figure 3, except that *K* = 1000 and that *p* and *θ* are variable. Here, extinctions can occur even when *p > p*^⋆^, because the population size is on the order of 1000, not infinite. Top panel: when plasticity and mutation rate are both low (dark region), the *A*-population goes extinct before a successful *B*-mutant arises. Bottom panel: when plasticity or mutation rate is large enough (dark region), the *A*-population is rescued (before extinction when *p* ≤ *p*^⋆^). Left: small rescue time (*T* ≤ 500); right: large rescue time (*T >* 500), infinite when *θ* = 0 and at the scale of genetic drift when *p* = 1.

- Increases the rate 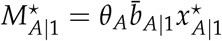 of mutant production;
- Decreases the probability 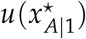 of mutant fixation.

Now the rate Λ of adaptation, seen as a function of phenotypic plasticity *p* on the interval [*p*^*⋆*^, 1], is a non-negative, continuous map vanishing at the extremities of the interval: Λ(*p*^*⋆*^) = 0 = Λ(1). Indeed, when *p* = *p*^*⋆*^, we have 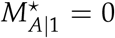 because 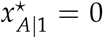 (recall we neglect here rescue events occurring on the path to equilibrium, here extinction). When *p* = 1, we have 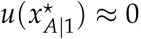, because each *B*-mutant is indistinguishable from the *A*-background, so that the probability of fixation is 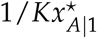, which vanishes when *K* is large.

Replacing 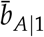 and *u* with their expression, we get (see Appendix)

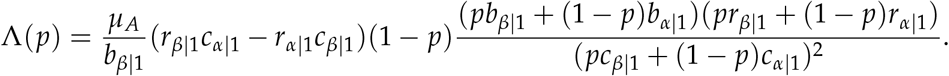

We can check directly on the formula that Λ(*p*^*⋆*^) = 0 = Λ(1). By Rolle’s Theorem, Λ performs its maximum at some *p*^*⋆⋆*^ ∈ (*p*^*⋆*^, 1), indeed justifying quantitatively that ‘intermediate levels of phenotypic plasticity favor evolutionary rescue’, as the article title claims (see Figure 5, last panel).

**Figure 5:**
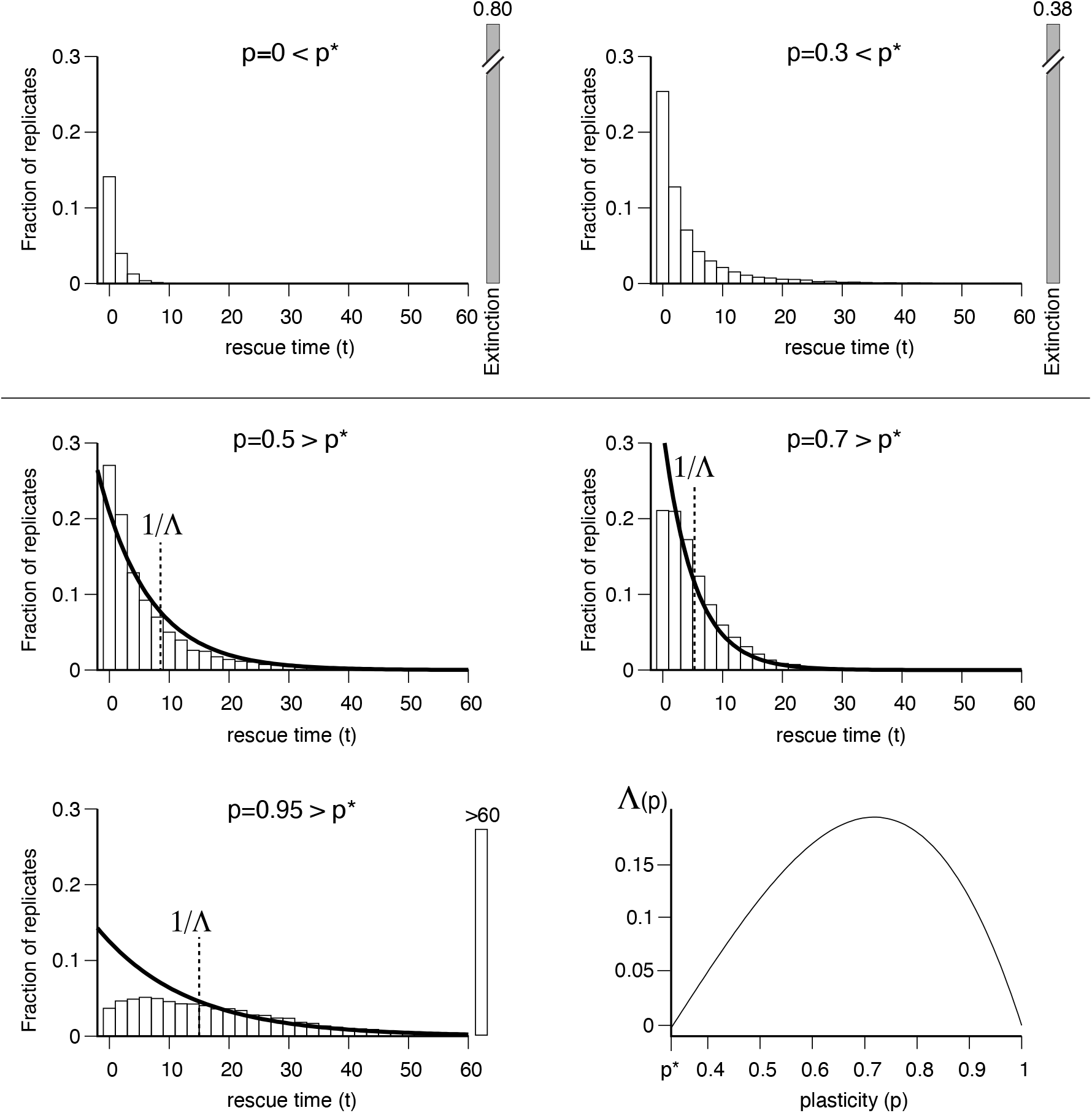
Rescue time *T* (waiting time to the emergence of a successful *B* mutant). In all panels, except bottom right, we report the empirical distribution (from 10^4^ replicates) of the rescue time (i.e., time elapsed between the change of environment and the appearance of a *B*-mutant that is not going extinct stochastically). Histogram bins have size 2. Parameters are identical to Figure 3, except *θ*_*A*_ = 1. The top panels are for *p < p*^⋆^, with an important fraction of extinctions. The medium and bottom left panels report results for *p > p*^⋆^, together with the expectation 1/Λ of rescue time and the discretized exponential distribution 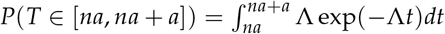, computed from Equation (6), where *a* denotes bin size. Bottom right: plot of rate of adaptation Λ(*p*) *vs p* ∈ [*p*^⋆^, 1] that shows a maximum in the vicinity of *p*^⋆⋆^ ∼ 0.7, which corresponds to a minimal rescue time.

Our results thus reconcile proponents of the Original Baldwin Effect and proponents of the Mayr Effect, in a way which depends on the level *p* of phenotypic plasticity:

- When *p < p*^*⋆*^, increasing *p* increases the probability of evolutionary rescue (i.e., arrival of an adaptive mutation before extinction) by delaying extinction. We call this effect the Strong Original Baldwin Effect, see Figure 5, top row;
- When *p* ∈ (*p*^*⋆*^, *p*^*⋆⋆*^), plasticity prevents extinction so that evolutionary rescue/adaptation (i.e., fixation of an adaptive mutation) occurs with probability 1 and increasing *p* even increases the rate of adaptation by increasing population size at equilibrium. We call this effect the Weak Original Baldwin Effect, see Figure 5, middle row;
- When *p > p*^*⋆⋆*^, evolutionary rescue occurs with probability 1, but increasing *p* reduces the fitness advantage of the adaptive mutants, thereby decreasing the rate of adaptation, i.e., delaying the time of effective rescue. This is the Mayr Effect, see Figure 5, bottom row.

## Discussion

### Theoretical implications

Our mathematical and numerical analyses have defined a series of distinct cases about the outcome of the population. The strength of plasticity conditions the survival of the plastic genotype A (A maintains when plasticity is above a threshold *p*^*⋆*^). In this ‘Survival’ area in Figure 6, the population can survive indefinitely, independently from the mutation rate. When the plasticity is maximal (*p* = 1), there is no more selection for B to replace A, and thus no expected genetic evolution beyond genetic drift (“Mayr Effect”, ME). The mutation rate from A to B conditions the probability that a genotype B appears and invades before the extinction of A. The ‘Rescue’ area in Figure 6 corresponds to cases where plasticity is not necessary to ensure population survival. The most interesting situation for which both plasticity and mutation alone are suboptimal, but allow for survival when combined, corresponds to a non trivial version of OBE (“Strong OBE”), when plasticity buys enough time for a rescue mutation to appear and invade the population.

**Figure 6:**
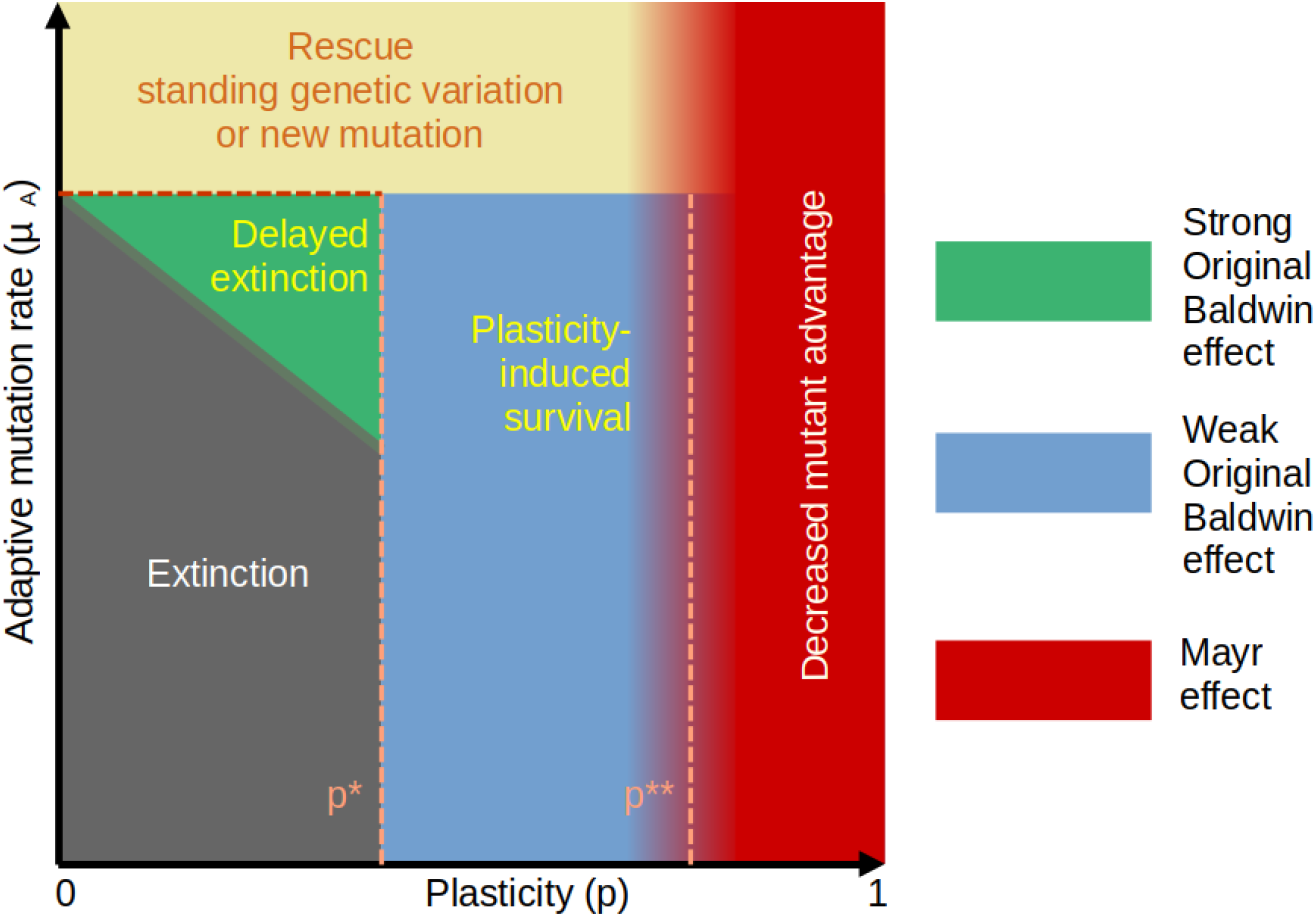
Summary representation of the different ways to survive (or not) environmental change in the adaptive mutation rate *vs* plasticity plane. For *p < p*^⋆^, the population goes to extinction unless a genetic variant arises (evolutionary rescue). In the green triangle, plasticity may delay enough extinction for rescue to occur despite moderate mutation rate (“Strong” OBE). When *p > p*^⋆^, plasticity sustains the population and increasing *p* has two antagonistic effects: to increase the supply of adaptive mutants by increasing population size (*p < p*^⋆⋆^, “Weak” OBE), and to decrease their probability of fixation by masking fitness differences (*p > p*^⋆⋆^, Mayr effect).

### Model limits

#### Noise vs plasticity

Here we chose to model plasticity by assuming that plastic genotypes respond deterministically in the phenotypic direction that maximizes fitness. However, our model remains relevant in situations where the phenotypic response to environmental change is not systematically adaptive. Noisy plastic responses would simply reduce the efficiency of the evolutionary rescue, by reducing the relative fitness advantage of plastic genotypes.

#### Cost of plasticity

In our theoretical framework, plasticity comes at no cost, i.e., a perfectly plastic genotype (p = 1) has the same fitness in the new environment as a genetically adapted variant. This remains highly debated (Auld et al. 2010; Murren et al. 2015). Even if plasticity was costly, the genetically adapted variant would fix more quickly in the population, but the model conclusions would remain roughly unchanged, except that the “drift” situation (neutral coexistence of both plastic and non-plastic variants) would not exist.

#### Periodic environments and the evolutionary origin of plasticity

Here, we have assumed the presence of a plastic genotype from the start, with no a priori hypothesis about the emergence of this plastic genotype. It is generally assumed that plasticity should realistically be maintained as a consequence of natural selection in fluctuating environments (Via and Lande 1985). Indeed, if in our model there is variation in the level of plasticity, our results suggest that the more plastic genotype will prevail by escaping extinction in the harsh environment *E*_1_ (at least in the absence of any cost of plasticity, see previous paragraph). If the adaptive mutation rate is not too small, the periodic alternation of environments will eventually lead, after a finite number of cycles, to the replacement of the plastic *A*-genotype by a mutant *B*-genotype adapted to *E*_1_. The population will then find itself in the reverse situation where *B* is the resident genotype and population survival depends on how plastic *B* is in the harsh environment *E*_0_. In such a ‘plasticity first’ setting, we thus expect the alternation of environments to drive the alternation of the dominant genotype, extending the motif studied here [(*Aα*|0)(*Aβ*|1)]^*n*^[(*Aα*|0)(*Bβ*|1)] to [(*Bβ*|1)(*Bα*|0)]^*n*^[(*Bβ*|1)(*Aα*|0)] and so on and so forth. In this periodic substitution sequence, each substitution, say from *A* to *B*, is facilitated by the partially plastic behavior of *A* in *E*_1_, which we denoted (*Aβ*|1), so that plasticity is constantly maintained, unless the population becomes extinct.

#### Plasticity-driven evolution

Here, the probability for an adaptive mutation to occur is the same in a highly-plastic *vs* a non-plastic genotype. This setting differs from Waddington’s intuition about genetic assimilation: as an embryologist, Waddington (1953*a*) thought that the same pathway would likely be involved first in the plastic response, and then in the final (‘assimilated’) genotype. In this view, recurrent selection of plastic phenotypes will lead to the accumulation of genetic variants up to a threshold where the adaptive phenotype will be constitutively expressed (Loison 2025). In this ‘plasticity first’ framework, plasticity is thought to pave the way to adaptive evolution by making genetically more accessible the genetic architecture that will ultimately enable the non-plastic expression of the adapted phenotype, a theoretical speculation that recently received partial piece of evidence. For instance, Levis et al. (2018) have shown that a new carnivorous morph of toad tadpoles have evolved from an ancestral herbivorous state through a series of plastic intermediates. There also exist examples where local adaptation was likely facilitated by fixing genetically some ancestral plastic states, such as for the clutch size in insects (Sun et al. 2020), and a case where an ancestral plastic genotype coexists in the wild with a derived non-plastic variant has recently been identified in Caenorhabditis (Vigne et al. 2021).

### Biological evidence of OBE

Countless examples of evolutionary rescue are now well-documented (Bell 2017). Evolutionary rescue can be studied in the lab through experimental evolution on microorganisms (Bell and Gonzalez 2009), and observed in the wild for a wide range of species, including plants (Bodbyl Roels and Kelly 2011) and vertebrates (Vander Wal et al. 2013). The vast majority of known cases of evolutionary rescue are associated with the undesirable emergence of resistant variants to antibiotics (Tazzyman and Bonhoeffer 2014), herbicides (Kreiner et al. 2018), or insecticides (Michaud 2018). The contribution of plasticity to evolutionary rescue has been studied theoretically (Chevin and Lande 2009; Chevin et al. 2010; Ashander et al. 2016), but empirical observations remain scarce (Chevin et al. 2013). Among several experimental issues, proving the effect of plasticity on the rescue probability requires to have both plastic and non-plastic genotypes available, which remains an exceptional situation. The case of persistent cells could be a potential candidate for a reproducible model system to study plasticity-facilitated evolutionary rescue. Drug-tolerant persisters are cells in tumors (McDonald and Dedhar 2024), fungi (Wuyts et al. 2018) or bacterial species (Fisher et al. 2017) that manage to switch plastically their physiology and stop dividing, thereby resisting a lethal drug concentration. The association between persistence and genetic resistance to antibiotics has been confirmed experimentally in E. coli (Levin-Reisman et al. 2017), in which mutations conferring resistance to ampicillin have been shown to occur preferentially in persister genetic backgrounds, rather that wild-type genotypes. The evolutionary scenario thus involves a plastic (persister) genotype delaying the antibiotic-related cell death, which opens the possibility for a beneficial mutation to occur and rescue the population. In spite of substantial differences with our theoretical setup (different reproduction and mutation rates among plastic and non-plastic genotypes), persister-related evolutionary rescue could thus qualify as a reproducible ‘strong’ Baldwin effect example.

The main consequence of the strong OBE is the unexpected possibility for populations to escape extinction, while being outside of the theoretical survival conditions by plasticity or by evolutionary rescue. This is particularly relevant when applying evolutionary models to conservation issues, especially in the context of the prediction of the species’ response to climate change, for which the interaction between evolutionary rescue and plasticity is a major determinant (Kelly 2019; Feiner et al. 2021). Understanding the factors involved in the probability of rescue in different environmental conditions is also crucial in microbiology and medical sciences (Lindsey et al. 2013), to avoid the emergence of genotypes resistant to antibiotics (Jalasvuori and Penttinen 2017) but also to understand the interactions between individuals and their microbiomes, which have been shown to contribute to the host resilience to stressful environments (Mueller et al. 2020).

### Historical and biological relevance of OBE

The question of whether Baldwin or Mayr was right can now be answered. For certain intermediate values of phenotypic plasticity, the intuition of Baldwin and the early proponents of “organic selection” takes the form of a non trivial joint of effect of mutation and plasticity on evolutionary rescue: our model allows to show that a strong version of OBE exists in a parametric region of its own, which does not overlap with a weaker version (where plasticity alone is enough for population survival). In this strong OBE situation, plasticity delays extinction of the population maladapted to the new environment, “buying time” for evolutionary rescue to happen.

In fact, it is when plasticity reaches a certain threshold that Mayr’s position becomes justified: genetic evolution by natural selection is first slowed down more and more sharply (OBE gets weaker and weaker, ME stronger and stronger), and then becomes the sole consequence of drift (ME). In this region of the parametric space, plasticity tends to erase too strongly the differences in fitness between genetic variants.

When exploring the parametric space on a large scale, the strong OBE region seems restricted, as it corresponds to a narrow combination of both adaptive mutation rate and level of plasticity. Yet, this does not necessarily mean that OBE is limited in natural conditions: both the mutation rate and the ancestral amount of plasticity are evolvable properties, and whether or not real species may lie close to this region of the parametric space remains to be determined, both theoretically and empirically. Contrary to what Simpson, Mayr and most of the founders of the Modern Synthesis thought, OBE may thus not be a negligible evolutionary process, especially when environments undergo significant alterations as a result of anthropogenic disturbances.

## Acknowledgements

The authors thank the two referees (R. Tucker Gilman and an anonymous referee) for their thorough review of a first version of the paper.

## Appendix Mathematical details

Recall that *N*_*i*_(*t*) denotes the number of individuals carrying allele *i* at time *t*.

First, we study the ecology in environment *E*_0_. Assume there is no *B*-individual initially. If the *A*-population starts in small numbers as in Figure 1, then because *r*_*α*|0_ *>* 0 (Assumption (1)), *N*_*A*_ grows like a supercritical branching process until it reaches a size of order *K* (unless it dies out rapidly, which happens with a probability strictly smaller than 1, even when *K* → ∞). At this point, if we write *X*_*A*_ = *N*_*A*_/*K*, the *per capita* birth rate is *b*_*α*|0_(1 − *θ*_*A*_/*K*) ≈ *b*_*α*|0_ and the *per capita* death rate is *d*_*α*|0_ + *c*_*α*|0_(*N*_*A*_ − 1)/*K* ≈ *d*_*α*|0_ + *c*_*α*|0_*X*_*A*_. Then as *K* → ∞, *X*_*A*_ converges to the solution of the following ODE

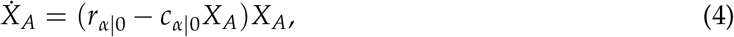

which indeed stabilizes around 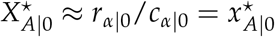, or equivalently 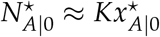.

In the main text, we have shown that at equilibrium, the *A*-population fuels the *B*-population by mutation at a global, constant rate *M* (which is formally the same as an immigration rate) and that every *B*-mutant start a branching process with *per capita* birth rate *B* and *per capita* death rate *D*, where *B < D*. This subcritical behavior is in line with our implicit assumption of neglecting the competition exerted by the descendants of mutants on the rest of the population (which prevents them from altering the equilibrium state of *N*_*A*_) as well as mutations from *B* to *A*. Then the (stochastic) equilibrium size of the *B*-population is given by the stationary distribution of the subcritical birth-death process with immigration, with immigration rate *M*, birth rate *B* and death rate *D*. This stationary distribution is given (Kendall 1948, 1949) by Equation (3).

At time 0, we assume that 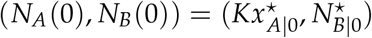, and the environment changes from state *E*_0_ to state *E*_1_. From then on, while *N*_*B*_ « *K, N*_*A*_/*K* behaves like the solution to the ODE

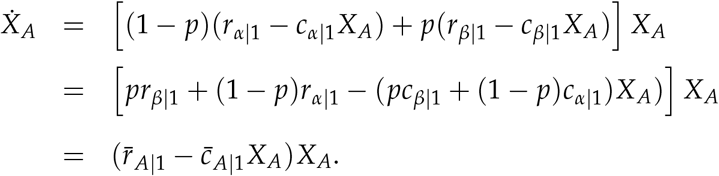

For *p* ≤ *p*^*⋆*^, we have 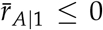 so that *X*_*A*_(*t*) decreases and converges to 0 as *t* → ∞, while for *p > p*^*⋆*^, we have 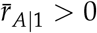 so that *X*_*A*_(*t*) converges as *t* → ∞, to

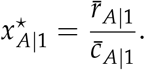

Also, for times *t* such that *N*_*B*_(*t*) remains negligible with respect to *K*, the *B*-population, formed of descendants of individuals present at time 0 and of *de novo* mutants, only feels the competition from the *A* population, which has approximately deterministic size *KX*_*A*_(*t*) at time *t*. Therefore, while *N*_*B*_ « *K*, the descendance size of any *B*-individual follows a time-inhomogeneous branching process with birth rate *b*_*β*|1_(1 − *θ*_*B*_/*K*) ≈ *b*_*β*|1_ and death rate at time *t d*_*β*|1_ + *c*_*β*|1_(*N*_*A*_(*t*) + *N*_*B*_(*t*) − 1)/*K* ≈ *d*_*β*|1_ + *c*_*β*|1_*X*_*A*_(*t*). We now inquire whether and when this time-inhomogeneous branching process can survive.

Provided *p* ∈ (*p*^*^, 1), we know that the equilibrium state of *X*_*A*_ in the absence of *B*’s is 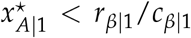. Then there is a time *t*^*⋆*^ such that for all *t > t*^*⋆*^, *X*_*A*_(*t*) *< r*_*β*|1_/*c*_*β*|1_ so that the growth rate of the *B*-population is positive. As a consequence, if we set

*v*(*t*) := *P* (the descendance of a single *B*-individual present at time *t* does not go extinct), then we have *v*(*t*) *>* 0 whenever *t > t*^*⋆*^.

Next, because the death rate of *B*’s present at time *t* + *s* depends on *X*_*A*_(*t* + *s*), *v*(*t*) depends on the whole trajectory (*X*_*A*_(*t* + *s*); *s* ≥ 0). More specifically, if for any positive function (*y*(*s*); *s* ≥ 0), we define *u*(*y*(*s*; *s* ≥ 0)) = *P* (the birth-death process started with one individual at time 0, with birth rate *b*_*β*|1_ and death rate *d*_*β*|1_ + *c*_*β*|1_*y*(*s*) at time *s*, does not go extinct), then we have

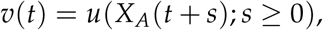

where it is implicit that 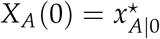 and is the solution to 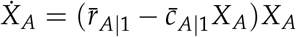. Now the probability that the descendance of a *B*-individual gets to size of order *K* (subsequently behaving deterministically) is approximately equal to the probability that this time-inhomogeneous branching process with positive growth rate escapes stochastic extinction. In other words, any *B*-carrier present at time *t* ≥ 0 has a probability of seeing its descendance get to a size of order *K* which converges to *v*(*t*) as *K* → ∞.

Unless we assume another change of environment (from *E*_1_ to *E*_0_), we are not interested in the dynamics following genetic adaptation, but let us describe briefly what happens once *N*_*B*_ is of order *K*. At this point, (*N*_*A*_/*K, N*_*B*_/*K*) is roughly equal the solution (*X*_*A*_, *X*_*B*_) to the following ODE

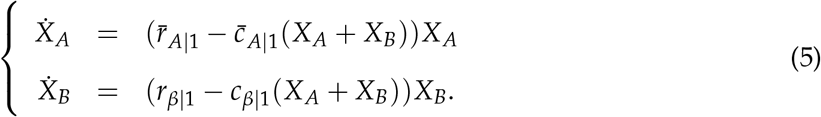

Now except when *p* = 1, we have 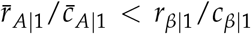 so the *B*’s outcompete the *A*’s, which end up in small numbers (following a subcritical branching process fueled by mutations from the *B*’s). When *p* = 1, *A*’s and *B*’s are indistinguishable, so that fixation can only occur by drift, which takes a time of order *K*.

We now come back to the rescue time *T*, defined as the time of appearance of the first *B*-mutant whose descendance does not become extinct. In particular, *T* = 0 if one of the preexisting *B*’s survives. Observe that each of them has the same probability *v*(0) to survive. Then by the branching property (holding so long as *N*_*B*_ « *K*), this probability converges as *K* → ∞ to

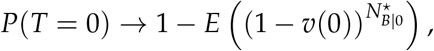

which can be computed thanks to Equation (3).

Now conditional on *T* /= 0, the rescue time *T* is larger than *t* iff the point process of mutation events giving rise to a surviving (i.e., not becoming extinct) *B*-population has no point in the interval [0, *t*]. As we showed previously, this point process is in the limit *K* → ∞ a time-inhomogeneous Poisson point process with rate 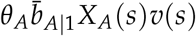 at time *s*, we get

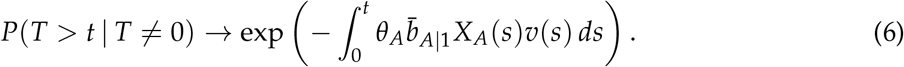

If we want to discard the possibility that adaptation proceeds from standing variation, we can take 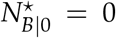 and focus on the previous formula to study how the rescue time *T* varies as a function of phenotypic plasticity *p*. In the main text, we focus on an approximation where *X*_*A*_ is constant (neglecting the transient phase until it reaches its equilibrium 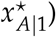, so that *T* is exponential and we show that *E*(*T*) has a minimum at some *p*^*⋆⋆*^ ∈ (*p*^*⋆*^, 1). Even when not making this approximation, we see from (6) that the variations of the law of *T* with *p* are non-monotonic, with *T* large both when *p* is small and *p* is close to 1:

- When *p < p*^*⋆*^, the function *X*_*A*_ is integrable in the absence of *B*-mutant (because as *t* → ∞, 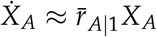 and 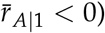, so that *P*(*T* = ∞) /= 0;
- As *p* → 1, the probability *v*(*s*) of non-extinction of a mutant appeared at time *s* goes to 0, so that *T* → ∞ in probability.

Let us now return to the approximation that when *p > p*^*⋆*^, *X*_*A*_ instantaneously reaches its new equilibrium 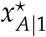 at time 0 (and also that 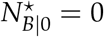, so that adaptation is not due to standing variation) to compute the rate of adaptation Λ = 1/*E*(*T*). Then 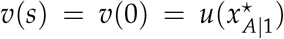 and according to (6), *T* follows the exponential distribution with parameter

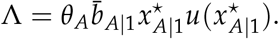

Since *P*(*T > t*) = *e*^−Λ*t*^, the dependence of *P*(*T > t*) upon *p* is the same for all *t* and studying how Λ varies with *p* gives the stochastic variation of *T* with *p*.

Recall that in the time-homogeneous case *u*(*x*) is the probability of non-extinction of a branching process with *per capita* birth rate *b*_*β*|1_ and *per capita* death rate *d*_*β*|1_ + *c*_*β*|1_*x*, which is equal to

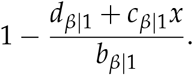

Replacing 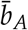 with its expression, we get

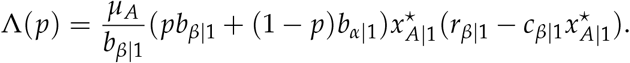

Also

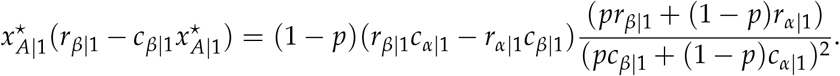

As a consequence, we get

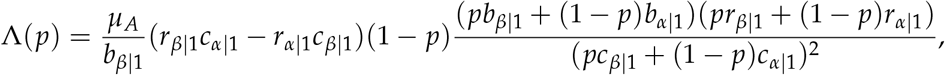

which is the formula given in the main text.

